# SmartImpute: A Targeted Imputation Framework for Single-cell Transcriptome Data

**DOI:** 10.1101/2024.07.15.603649

**Authors:** Sijie Yao, Xiaoqing Yu, Xuefeng Wang

## Abstract

Single-cell RNA sequencing (scRNA-seq) has revolutionized our understanding of cellular heterogeneity and tissue transcriptomic complexity. However, the high frequency of dropout events in scRNA-seq data complicates downstream analyses such as cell type identification and trajectory inference. Existing imputation methods address the dropout problem but face limitations such as high computational cost and risk of over-imputation. We present SmartImpute, a novel computational framework designed for targeted imputation of scRNA-seq data. SmartImpute focuses on a predefined set of marker genes, enhancing the biological relevance and computational efficiency of the imputation process while minimizing the risk of model misspecification. Utilizing a modified Generative Adversarial Imputation Network architecture, SmartImpute accurately imputes the missing gene expression and distinguishes between true biological zeros and missing values, preventing overfitting and preserving biologically relevant zeros. To ensure reproducibility, we also provide a function based on the GPT4 model to create target gene panels depending on the tissue types and research context. Our results, based on scRNA-seq data from head and neck squamous cell carcinoma and human bone marrow, demonstrate that SmartImpute significantly enhances cell type annotation and clustering accuracy while reducing computational burden. Benchmarking against other imputation methods highlights SmartImpute’s superior performance in terms of both accuracy and efficiency. Overall, SmartImpute provides a lightweight, efficient, and biologically relevant solution for addressing dropout events in scRNA-seq data, facilitating deeper insights into cellular heterogeneity and disease progression. Furthermore, SmartImpute’s targeted approach can be extended to spatial omics data, which also contain many missing values.

## INTRODUCTION

Over the past decade, the quick development of single-cell RNA sequencing (scRNA-seq) has revolutionized our understanding of cellular heterogeneity and tissue transcriptomic complexity. By delivering high-resolution gene expression data at the individual cell level, scRNA-seq facilitates the in-depth study of cellular subtypes, lineage relationships, and dynamic biological processes with unparalleled detail. However, a significant challenge in scRNA-seq data is the high frequency of dropout events, where gene expression measurements are recorded as zero due to several reasons: (1) low amounts of mRNA in individual cells, (2) technical or sequencing artifacts, and (3) inherent cell type differences, with some cells exhibiting low expression levels for certain genes. These dropouts complicate downstream analyses, such as cell type identification ^1,2^, differential expression analysis, and trajectory inference ^3-5^, thereby necessitating imputation methods to infer the missing values before proceeding with downstream scRNA-seq analysis.

Existing imputation methods, such as SAVER ^6^, SAVER-X ^7^, DCA ^8^, and DeepImpute ^9^, have made significant progress in addressing the dropout problem. SAVER uses a Bayesian framework, assuming a Poisson-Gamma mixture model to estimate gene expression levels, while DCA employs a zero-inflated negative binomial noise model within an autoencoder framework ^10,11^. DeepImpute, on the other hand, utilizes a deep neural network approach to impute target genes. SAVER-X combines a deep autoencoder with a Bayesian model and extracts transferable gene-gene relationships across different datasets and even different species to denoise new target datasets. Despite their advancements, these methods face several limitations that restrict their applicability in daily RNA-seq or large-scale studies. They often impose a high computational cost, risk overfitting and producing biased results, and require users to manage larger imputed matrices compared to sparse ones. Additionally, in many real data analyses, imputation may offer limited improvement in downstream analyses such as cell typing. These issues underscore the need for more efficient and biologically relevant imputation methods to address the challenges posed by dropout events in scRNA-seq data.

In this study, we present SmartImpute, a novel computational framework specifically designed for imputing targeted scRNA-seq data. SmartImpute focuses on a predefined set of marker genes most informative for understanding cellular behavior. This approach is based on observations in cancer genomics data analysis, where despite the heterogeneity in both cancer and immune cells, most cell types and subtypes can be identified using fewer than 1000 genes. This targeted strategy not only improves the biological relevance of the imputation process but also enhances computational efficiency by limiting the imputation to the most informative subset of genes. Moreover, SmartImpute leverages a modified Generative Adversarial Network (GAN) architecture ^12^, known as GAIN ^13^, to accurately distinguish between true biological zeros and missing values. By integrating a multi-task discriminator ^14,15^, SmartImpute prevents overfitting and maintains the integrity of biologically relevant zeros, thereby ensuring high imputation accuracy.

We demonstrate the performance and efficiency of SmartImpute using scRNA-seq data from head and neck squamous cell carcinoma (HNSCC) and human bone marrow. Our results show that SmartImpute significantly enhances the visualization of cell type clusters, improves cell type annotation, and preserves biologically meaningful gene expression patterns. Benchmarking against other imputation methods highlights SmartImpute’s superior performance in terms of both accuracy and computational efficiency. Overall, SmartImpute provides a lightweight and more realistic solution for addressing dropout events in scRNA-seq data. By focusing on targeted gene panels and employing an advanced GAN architecture, SmartImpute enhances the biological interpretability of single-cell data, facilitating deeper insights into cellular heterogeneity and disease progression.

## RESULTS

### Targeted marker gene panel approach

SmartImpute introduces a targeted approach to the imputation of scRNA-seq data by selectively focusing on a predefined set of marker genes. Using a targeted gene panel offers two key benefits. First, it improves the biological relevance of the imputation process by concentrating on genes important for understanding cellular behavior. Second, it makes the computation more efficient by only imputing on the most informative gene subset of the data. Starting with a group of 580 well-established marker genes (BD Rhapsody™ Immune Response Targeted Panel (Human)), researchers have the flexibility to adjust a customized panel to fit the needs of their specific projects. This customization enabled a tailored imputation that not only aligns with specific research objectives but also embeds biological insights directly into the imputation process, setting it apart from methods such as DeepImpute that may overlook biological significance. To aid in the selection of an effective gene panel with specific information about data, we offer an R package that utilizes a Generative Pre-trained Transformer (GPT) model integrated with SmartImpute framework. (**Star Methods**).

### Modified GAN Architecture for Imputation

SmartImpute employs a modified version of Generative Adversarial Nets (GAN) (**Star Methods**) to address the imputation challenge in scRNA-seq. The process begins with data preparation by identifying the target gene panel in the gene expression data. The training process is refined by incorporating a proportion of non-target genes, without losing information from the dataset and ensuring robust model generalizability. SmartImpute uses a multi-task discriminator to improve the imputation accuracy and the proper identification of biological zeros (**Figure. 1**). This modified generative model allows SmartImpute to impute values without making any assumptions about the underlying data distribution. Consequently, the imputed values produced by SmartImpute exhibit greater accuracy compared to other methods such as SAVER, which relies on a Poisson-Gamma mixture model, and DCA, which utilizes an autoencoder (AE) model.

**Figure 1.**
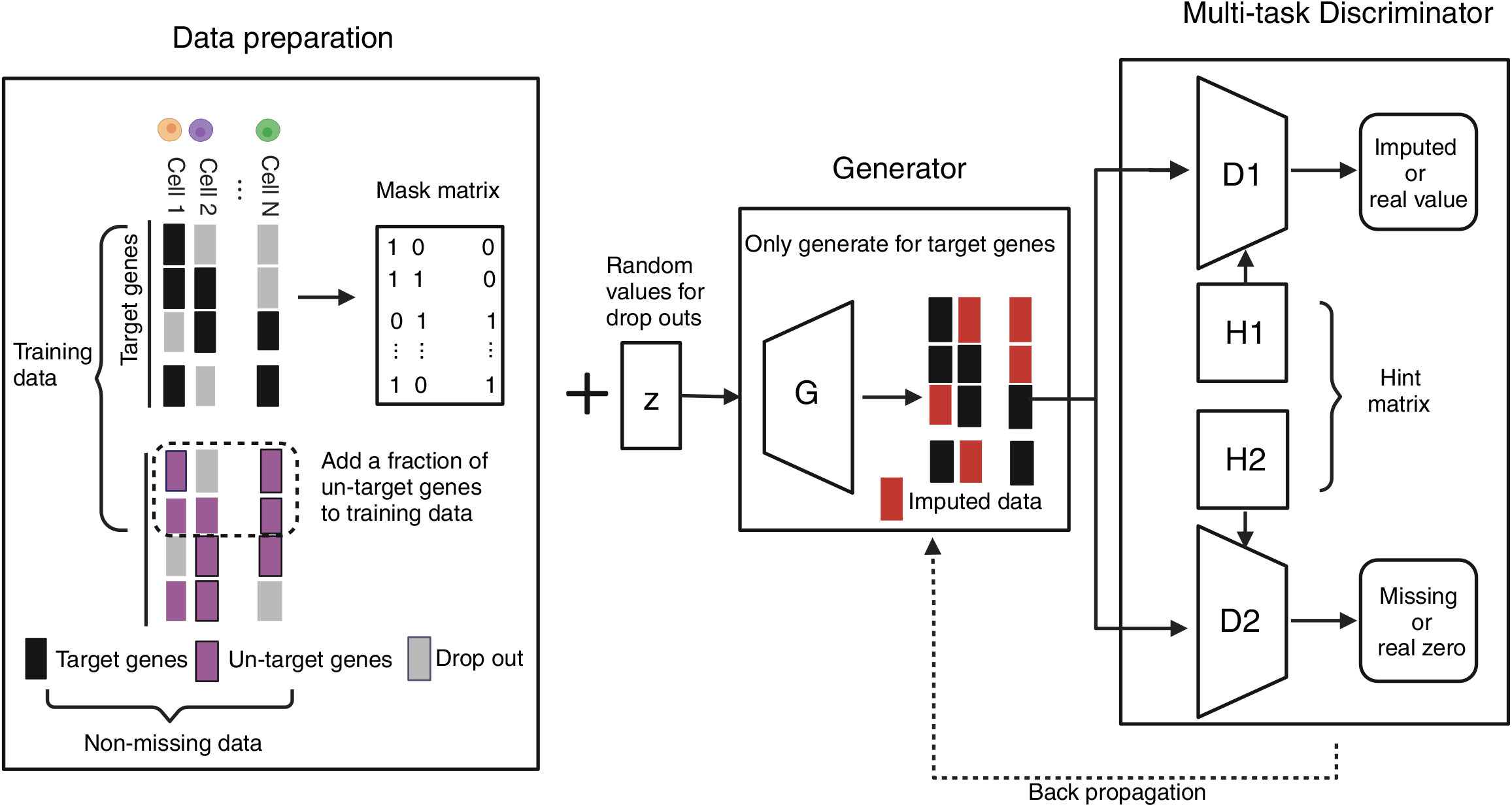
Overview of SmartImpute: SmartImpute uses a targeted approach for imputing missing value in scRNA-seq data. It adapts the GAIN architecture, adding a multi-task discriminator to distinguish between biological zeros and missing values.

### Performance of SmartImpute in HNSCC Normalized Data

To evaluate SmartImpute’s performance in normalized data, we conducted a comparison of UMAP clustering using head and neck squamous cell carcinoma (HNSCC) data with and without imputation. The visual comparison demonstrates SmartImpute’s ability to distinguish between different cell types, which is not feasible without imputation. Particularly, cell types such as CD4 Tconv, CD8 exhausted, and CD8 Tconv are indistinguishable without imputation. Moreover, SmartImpute is able to separate closely related cell types like Myocytes, Fibroblasts, and Myofibroblasts while preserving their biological distinctiveness (**Figure. 2A**).

**Figure 2.**
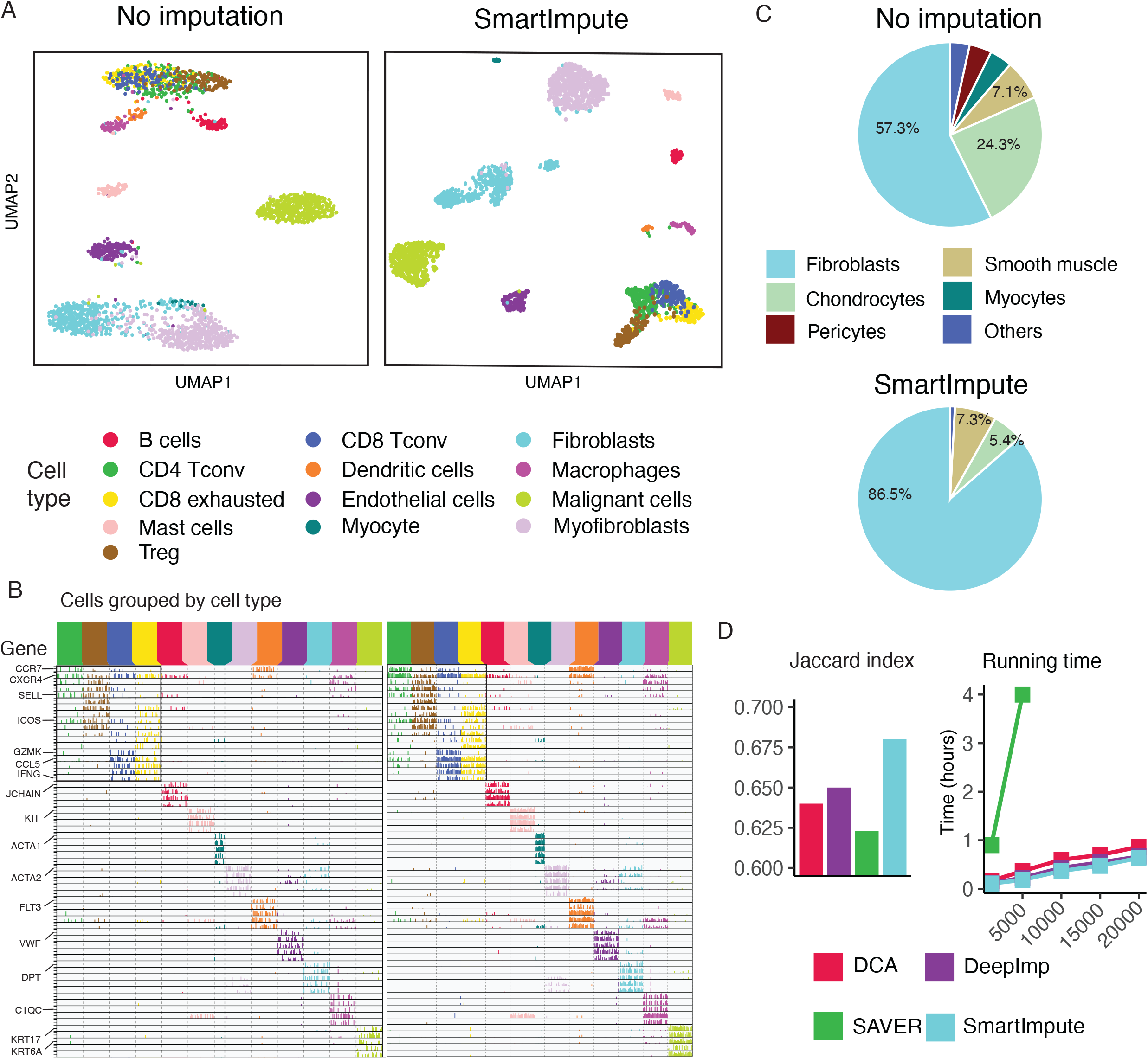
Performance of SmartImpute in HNSCC normalized data: (A) UMAP clustering without imputation (left) and with SmartImpute (right) (B) Gene expression heatmap for cell type associated marker genes without imputation (left) and with SmartImpute (right) (C) SingleR cell type annotation accuracy in Fibroblasts (D) Comparison with different imputation methods: Jaccard index (left) and running time (right).

Next, we demonstrate the accuracy of SmartImpute’s imputation by comparing heatmaps of gene expressions with and without imputation. Ideally, the heatmap should present distinct, filled diagonal blocks, each representing the expression of marker genes for specific cell types. For off-diagonal blocks, they should remain empty since the genes are not expressed in the corresponding cell types. In the heatmap without imputation, the block pattern can be observed for marker gene associated cell types. However, the pattern is weak caused by the great amount of dropout events inside these blocks, leading to a sparse expression structure and making downstream analysis more difficult. In contrast, the heatmap after applying SmartImpute shows significant improvement. The imputation process fills in the missing values, leading to more complete and continuous expression patterns across all cell types. Specifically, the marker gene blocks are distinctly filled, particularly for T cells (CD4 Tconv, CD8 exhausted, and CD8 Tconv), showcasing clearly defined clusters of gene expression. Genes important for specific cell types, such as CCR7, CXCR4, and ICOS for immune cells, show more consistent and biologically relevant expression patterns with imputation. Moreover, the sparsity of data in the off-diagonal blocks remains after imputation, indicating that SmartImpute prevents the overfitting without generating false positives (**Figure 2B**).

To check the performance with SmartImpute for downstream analysis, we employed SingleR ^16^ with a BLUEPRINT cell type reference, containing most cell types present in this dataset, to annotate cell types with and without imputation. With imputation, all the annotation results are better than without imputation, and this improvement is particularly observed in Fibroblasts. SingleR identifies only 57.3% of true Fibroblasts without imputation, whereas SmartImpute increases this accuracy to 86.5% (**Figure. 2C**). Additional results highlighting improvements in cell typing across other cell types are provided in **Supplementary Figure 1A**. Similarly, another cell typing method, AUCell ^17^, shows SmartImpute’s ability to enhance cell typing accuracy, particularly with malignant cells (**Supplementary Figure 1B)**.

To address the advantage of targeted imputation approach from SmartImpute, we benchmarked SmartImpute against other imputation methods from accuracy and efficiency. In the accuracy comparison, SmartImpute achieves the highest Jaccard Index, with a score of 0.678, outperforming DCA (0.637), DeepImp (0.649), and SAVER (0.623) (**Figure 2D)**. The UMAP clustering for these scRNA-seq imputation methods is given in **Supplementary Figure 2**. To illustrate SmartImpute’s computation efficiency, we compare the running times with an increasing number of genes. SAVER is unable to process datasets nwith more than 5000 genes, while SmartImpute consistently maintains the shortest running time in comparison to the other methods (**Figure 2D**).

### Performance of SmartImpute in HNSCC UMI count data

Next, we evaluate SmartImpute’s performance under a more complicated scenario using Unique Molecular Identifier (UMI) count data. Normalization often reduces biases such as sequencing depth, batch effects, and other technical artifacts. Consequently, imputing raw count data is more difficult compared to normalized data. We use UMI count data from HNSCC to check the performance of SmartImpute.

Similar to the results in normalized data, we show SmartImpute’s efficacy in resolving different cell types into coherent clusters by UMAP clustering (**Figure 3A**). Our focus remains on the CD8 T cell compartment and its difference before and after imputation. The CD8 T cell subtypes—CD8 TExh (exhausted), CD8 TpreExh (pre-exhausted), and CD8 TEM (effector memory)—play a pivotal role in immune response modulation and exhibit close phenotypic and functional relationships. In the PBMC reference, these subtypes are collectively identified as CD8 TEM (PAN) cells due to their shared effector functions and memory characteristics. Proper cell typing should annotate these subtypes under the broader category of CD8 TEM (PAN) when referenced with PBMC data. The cell typing accuracy is visualized in Sankey diagram, where the improvement of cell type identification is observed after imputation (**Figure 3C**). Without imputation, a significant number of CD8 TExh cells are misclassified as CD8 Naïve. Additionally, about one-fourth of CD8 TpreExh and CD8 TEM cells are incorrectly labeled as CD4, CD8 TCM, or other T cells. After imputation, these misclassifications are significantly reduced. Almost no CD8 TExh cells are annotated as CD8 Naïve, and misclassifications to other T cells from CD8 TExh, CD8 TpreExh and CD8 TEM are reduced to one-third of the original rate. To quantitively measure this improvement, we calculate the F1 score for cell typing accuracy with and without imputation. SmartImpute improves the F1 score from 0.61 to 0.73, for CD8 TpreExh from 0.63 to 0.71, and for CD8 TEM from 0.62 to 0.75. Notably, other cell types also show a general improvement in cell typing accuracy, with the F1 score increasing from 0.71 to 0.75 after imputation (**Figure 3D**).

**Figure 3.**
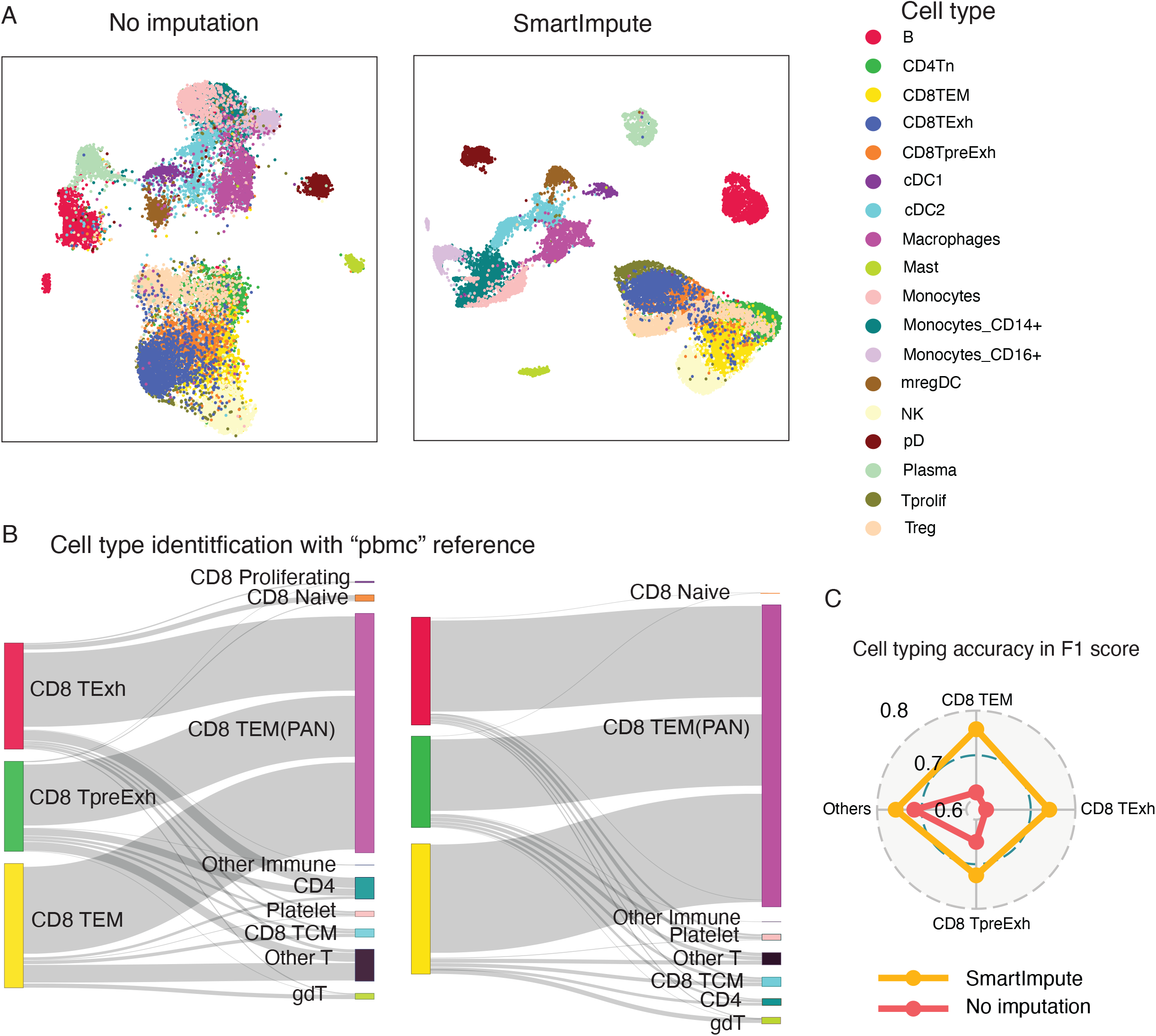
Performance of SmartImpute in HNSCC UMI count data: (A) UMAP clustering without imputation (left) and with SmartImpute (right) (B) Cell type identification for CD8 T cells without imputation (left) and with SmartImpute (right) (C) Cell typing accuracy measured in F1 score.

### SmartImpute improves Classification in T cell subtypes

Furthermore, we apply SmartImpute on an even more challenging problem: separating closely related T-cell subtypes. These subtypes are biologically similar, making it difficult to distinguish them in downstream analyses. We demonstrate the capability of SmartImpute in effectively separating different T cell subtypes using T cell data. The clarity in cluster distinction among various T cell populations, is significantly improved when SmartImpute is applied. This is observed in the separation of cell type clusters such as CD4T naïve and Th2, with the imputed data demonstrating a higher degree of cluster cohesiveness and a reduction in misclassifications (**Supplementary Figure 3A**). Quantitative improvements are measured in T cell subtype clustering results, with Jaccard Index (JI) increasing from 0.48 to 0.63 and Adjusted Rand Index (ARI) increasing from 0.46 to 0.65 after imputation with SmartImpute (**Supplementary Figure 3B**).

### Unraveling Cellular Trajectories with SmartImpute in Human Bone Marrow Data

The targeted imputation approach in SmartImpute is further highlighted in a BM single cell data using a custom gene panel with 462 genes (BD Rhapsody™ Immune Response Targeted Panel (Human)). The original study utilized both mRNA and protein surface markers to facilitate cell clustering and trajectory analysis. In contrast, our aim is to reach equivalent levels of downstream analysis performance only using mRNA data imputed by SmartImpute. After imputation, the UMAP clustering plot clearly shows how Hematopoietic Stem Cells (HSCs) and Multipotent Progenitors differentiate into various cell lineages. This includes the paths from HSCs to blood cells, B cells, and natural killer (NK) cell progenitors. Additionally, the progression of Pro B cells into more mature B cell stages is clearly observed. Moreover, the imputed data reveals how Monocytes differentiate into various types of Dendritic Cells, presenting a comprehensive picture of immune cell development. (**Supplementary Figure 4**). This downstream analysis for cell trajectories demonstrates SmartImpute’s ability to find complex cellular lineage pathways with only a target mRNA panel, achieving results that are similar with the original study that leverages additional information of protein surface markers.

Collectively, our evaluation of SmartImpute’s performance reveals its capability in imputing scRNA-seq data across various datasets. By customizing the gene panel selection process and integrating GAIN architecture, SmartImpute consistently delivers highly accurate results that have significant implications for the interpretation and analysis of single-cell data.

## DISCUSSION

The targeted imputation approach employed by the proposed SmartImpute offers significant advantages in enhancing both the biological relevance and computational efficiency of scRNA-seq data analysis. By focusing on a predefined set of marker genes, SmartImpute aligns the imputation process with specific biological contexts, ensuring that the imputed values are meaningful for understanding cellular behavior. This strategy is particularly beneficial for studies where certain genes are known to be critical for distinguishing between cell types, subtypes, or states. For example, in the single-cell sequencing datasets evaluated in this study, SmartImpute exhibited superior or comparable performance in improving cell type annotation and clustering accuracy relative to other imputation methods. The use of a targeted gene panel not only improves the interpretability of the results but also significantly reduces the computational burden, allowing for faster processing and scalability to larger datasets. Another benefit of this approach is that, since only a subset of genes is imputed, the risk of overfitting is minimized, and data integrity is better preserved compared to full-panel imputation. Overall, SmartImpute has the potential to be a lightweight and effective tool for routine single-cell analysis.

The targeted imputation approach can also be implemented using other computational frameworks besides GAIN. For example, generative models based on Variational Autoencoders (VAEs) could be modified to achieve targeted imputation. VAEs are effective in capturing the underlying data distribution and generating less biased imputations by learning a latent space representation. Compared to GAIN, VAEs typically offer more stable training and are less prone to mode collapse and convergence failure. However, GANs often produce more accurate imputations due to their adversarial training process, which encourages the generation of high-resolution gene expression profiles. The choice between GAIN and VAEs for targeted imputation would depend on the specific requirements of the study, such as the need for stable training (favoring VAE) or higher accuracy in imputed values (favoring GAN). Implementing targeted imputation with different generative models, or even in an integrated framework, could further expand the flexibility and applicability of this approach in various single-cell RNA sequencing analyses. Both frameworks also have the potential to benefit from pre-training models based on other datasets, facilitating transfer learning similar to the approach implemented in SAVER-X. This would enable the models to leverage existing biological knowledge and improve imputation accuracy across different datasets and conditions.

Despite its promising strengths, the targeted approach also has limitations that must be considered. One potential drawback is the reliance on prior biological knowledge to select the marker genes, which may introduce biases if the selected panel does not capture all relevant genes for a given study. This could result in the omission of important biological signals and potentially impact downstream analyses. Moreover, the predefined gene panels may need to be frequently updated to reflect new discoveries in the field, adding an additional layer of complexity to the imputation process. To encourage reproducibility, we have provided a GPT model for automatic gene panel selection, which partially addresses the issue by enabling the customization of gene panels. However, it is essential to validate the selected genes rigorously to ensure their relevance and comprehensiveness for specific research context and experimental designs. Additionally, it is important to conduct further research to identify more informative biomarkers or panels for different cell types or cell states, thereby enhancing the overall effectiveness and applicability of targeted imputation in various single-cell RNA sequencing studies.

The same targeted imputation method can also be extended to spatial omics data, such as NanoString CosMx and 10X Xenium, which often contain many missing values. It is important to note that many spatial omics panels are targeted by nature, and with larger panels, the proportion of missing values tends to increase. By applying SmartImpute’s targeted approach to spatial transcriptomics, researchers can achieve similar improvements in data quality and interpretability, facilitating deeper insights into the molecular mechanisms associated with the spatial organization of tissues.

## STAR★Methods

### METHOD DETAILS

#### SmartImpute overview

SmartImpute employs a targeted imputation approach for single-cell RNA sequencing data, using a select panel of marker genes essential for identifying distinct cell types and states. This approach leverages prior biological knowledge to enhance imputation accuracy and computational efficiency. SmartImpute adapts a modified Generative Adversarial Network (GAN) architecture, specifically tailored for imputation tasks, known as GAIN. The original GAIN framework is limited in scRNA-seq applications where dropouts are represented as zero values, lacking the capability to differentiate between biological zeros and missing data. To address this, SmartImpute incorporates an additional discriminator aiming for distinguishing between the biological zeros and missing values in scRNA-seq data. In the following, we will discuss (1) the implementation of target imputation approach, (2) the multi-task discriminator for imputation in scRNA-seq data (3) target panel genes selection, (4) training details and (5) evaluation metrics.

#### Targeted imputation with attention mechanism

SmartImpute improves accuracy and computational efficiency by incorporating biological information through its targeted imputation approach. This approach is implemented in the generator (G) by incorporating an attention mechanism. For clarity, the following notation is introduced: the expression matrix of single cell data is denoted by a P by N matrix that *X* = (***x***_1_,…, ***x***_*N*_), ***x***_*i*_ = (*x*_*i*1_,., *x*_*iP*_)′. Without losing generality, the first q genes are designated as the target genes. In the targeted imputation setting, only dropouts in ***x***_*i*_ where *i* ≤ *q* will be imputed.

An attention parameter *γ* is incorporated to refine the generating process. The attention parameter *γ* takes range between 0 and 1, allowing for the integration of additional genes from outside the target panel to aid in the generation of missing values. During model training, each batch is composed of target genes ***x***_1_ to ***x***_*q*_, and a supplementary set of [*γ* ⋅ (*p* - *q*)] randomly chosen genes with [·] denoted as the ceiling function. In SmartImpute, the generator G consists of K subnets, denoted as *G*_*k*_, *k* = 1,…, *K*. During the training of the k-th batch, the subnet *G*_*k*_ functions by taking the expression matrix X, random noise Z, and attention parameter *γ* as inputs:

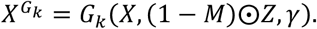

Here M is the mask matrix which indicating the dropouts event with: *m*_*ij*_ = 1 for *x*_*ij*_ ≠ 0 and *m*_*ij*_ = 0 otherwise. The term (1 - *M*)⨀*Z* ensures that the generator only incorporates random noise corresponding to the positions of dropouts during the training process. The final output of generator G is the average of all subnets output 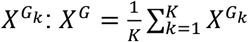.

#### Biological zero recognition with multi-task discriminator

Although the target imputation approach already considers the issue of overfitting, SmartImpute further prevent it by adding a multi-task discriminator. Some Imputation methods such as DCA and DeepImpute consider every dropout as a missing value and impute expression values for all genes in every cell. While SAVER applies a lasso penalization to preserve biological zeros but identifies relatively few of them. To address this challenge, we introduce a biological zero recognition task with SmartImpute using a multitask discriminator.

The multi-task discriminator is designed to handle two tasks. The first discriminator *D*_1_performs the same function to the original discriminator in the GAIN framework. It identifies whether elements of *X*^*G*^ are real or imputed. To accomplish the task, *D*_1_ takes the mask matrix M of original data as one of its inputs. The primary objective of *D*_1_ is to accurately reproduce the mask matrix M from the imputed dataset *X*^*G*^. While *D*_1_ can distinguish between real and imputed data, it will impute expression values of target genes of non-related cell types. To address this issue and prevent the overfitting of imputations on biological zeros, an additional discriminator *D*_2_ is incorporated in the framework.

The second discriminator *D*_2_, addresses a unique challenge in scRNA-seq data by determining whether the observed dropouts represent true zeros, indicating no gene expression, or missing values due to technical limitations. Different to *D*_1_, *D*_2_ employs a specifically designed missing assignment matrix 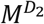 to allocate missing zeros for the imputed data with 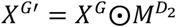. To enhance the identification of biological zeros, *D*_2_ incorporates a threshold value *τ* which is used to classify gene expression values. When the expression values are below the threshold *τ*, these elements are considered as biological zeros: 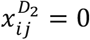 if 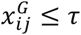. Otherwise, it keeps its value from 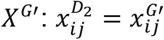. This approach ensures that 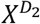contains the true biological zeros information even when there are no zeros in the imputed data *X*^*G*^. The mask matrix for biological zero recognition 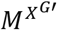 is further updated with 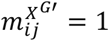 if 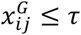, and 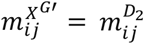, otherwise.

#### Double hint mechanism

Since two discriminators *D*_1_ and *D*_2_ makes discriminations on different input data, a double hint mechanism is used in SmartImpute framework. Following the hint mechanism in GAIN, each discriminator in SmartImpute will be assigned with distinct hint matrices. By providing partial information about the missing entries, the hint mechanism guides the discriminator towards a more well-defined and unique optimal solution. This refinement in the optimization landscape enhances the robustness by the convergence of training process and subsequently improves the quality of the imputations.

For *D*_1_, the hint matrix H1 is derived from the mask matrix of the original data. *H*_1_ provides part of information by indicating which values in the data matrix are observed versus those that are missing:

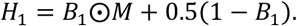

Where *B*_1_ is a random variable taking the value from ^16^. The hint matrix is used as one of the inputs for discriminator, and *D*_1_ is constructed as a function of *X*^*G*^, *H*_1_ and 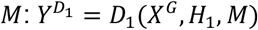.

For the discriminator *D*_2_, the corresponding hint matrix *H*_2_ is defined by 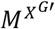,

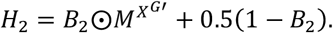

*B*_2_ performs the same function as *B*_1_ for a random assignment of where to give hint and *D*_2_ is the function of *X*^*G*′^, *H*_2_ and 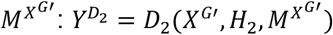.

#### Optimizing imputation with adversarial training by WGAN

SmartImpute takes the advantage of GAN model in improving the imputation accuracy with an adversarial training process. SmartImpute’s training procedure involves an adversarial training strategy, where the generator competes with the discriminators to improve imputation accuracy. The imputation model of SmartImpute is iteratively trained by fixing one of the generator or discriminator at a time with back propagation. For the training of generator, two loss function is used:

1. Adversarial Loss (Wasserstein Distance): The first loss function of generator reflects the adversarial process of GAN architecture. In the adversarial process, the generator is designed to fool the discriminator *D*_1_ by minimizing this loss, while Discriminator *D*_1_ aims to accurately differentiate between the data from generator and the original data by maximizing the same function. This adversarial loss is measured using the Wasserstein distance, known as WGAN ^18^. WGAN’s advantages include more stable training, meaningful and interpretable loss metrics, reduced mode collapse, robustness to hyperparameters, and ultimately higher-quality generated samples. The loss function with Wasserstein distance is defined as:

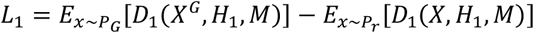

In this context, *P*_*r*_ and *P*_*G*_ denote the real and generated data distributions, respectively.
2. Reconstruction Loss: The second part of the generator’s loss function focuses on the accuracy of the imputed data. The generated values for the observed elements should be close to the original values. This objective is defined using the mean squared error (MSE) for normalized (continuous) data:

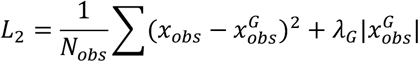

The reconstruction loss is calculated only with the observed entries. Here, *N*_*obs*_ is the number of observed entries, *x*_*obs*_ are the original observed values, and 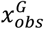are the generated values. We add an extra regularization term 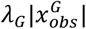 to threshold the biological zeros.

When dealing with the UMI count data, SmartImpute adapts the reconstruction loss by a zero-inflated negative binomial (ZINB) likelihood:

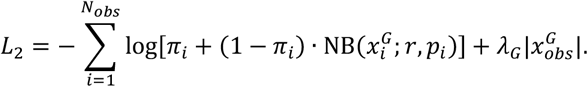

Here *π*_*i*_ is the probability of an excess zero for the i-th gene, and 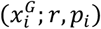 is the probability mass function of the negative binomial distribution with success probability *p*_*i*_ and number of failures r before the i-th success.

The loss function for generator is then the weighted sum of *L*_1_ and *L*_2_:

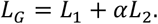

For discriminator *D*_1_, the loss function is the negative of *L*_1_,

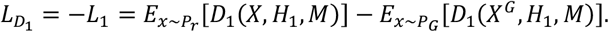

And for discriminator *D*_2_, we train *D*_2_ to maximize the probability of *D*_2_ predicting *M*^*G*′^,

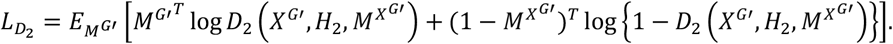

Where log is element-wise logarithm, and the expectation is calculated based on the random variable*M*^*G*′^.

#### Hyperparameter tuning

We conducted a hyperparameter search for different datasets using SmartImpute. To maintain robustness and computational efficiency, we fixed as many hyperparameters as possible. The main hyperparameters to search in SmartImpute includes the attention parameter *γ*, biological zero regularization parameter *λ*_*G*_, hint rate for *H*_1_: *h*_1_, hint rate for *H*_2_: *h*_2_, learning rate *ε*_*E*_, and weighting parameter *α*.

For the normalized HNSCC data, we first performed a 5-fold cross-validation to train the multi-task targeted GAIN. Two critical parameters were decided with *ε*_*E*_ = 0.05 and *λ*_*G*_ = 0.5. In this dataset, we set the number of hidden layers to 2 with nodes number 64. For SmartImpute, we found that three hidden layers for G and two for both *D*_1_ and *D*_2_ gave the relatively robust results with the number of nodes: 128, 128, 128, 32, and 32. Results were produced with learning late 0.001, *h*_1_=*h*_2_ =0.9, attention parameter *γ* 0.005, number of subnets 4, number of sample batch 64, penalty *λ*_*G*_=1, and weighting parameter *α* = 90.

For the HNSCC count data, we train the generator with ZINB likelihood function for *L*_2_. The hyper-parameters values were given with with learning late 0.001, *h*_1_=*h*_2_ =0.8, attention parameter *γ* 0.002, number of subnets 4, number of sample batch 64, penalty *λ*_*G*_=1, and weighting parameter *α* = 75.

#### GPT model for Target Gene Panel Selection

To optimize the targeted imputation approach employed by SmartImpute, the selection of an informative gene panel is performed using a GPT model with the R software package tpGPT. A well-established default base gene panel of 580 marker genes (BD Rhapsody™ Immune Response Targeted Panel (Human)). is provided as a starting point for constructing a data-specific target panel. In tpGPT, users can input specific parameters such as cancer type, tissue type, and cell type to add extra genes to the default panel for their own studies. The study-based gene list is generated using a structured prompt message based on the following template:

‘‘Generate a list to Number marker genes in Cancer_type associating with Tissue_type. \n Only provide the gene list. \n Ensure each gene is listed in a new line without any prefixes such as numbers.”

In this template, “Number”, “Cancer_type” and “Tissue_type” are placeholders that will be replaced with the corresponding parameters. (For example, 250 for “Number”, “head and neck cancer” for “Cancer_type” and “epithelial tissue” for Tissue type.)

For generating a cell type-specific gene panel, the following prompt template is used:

“Generate a list of marker genes for following cell types: Cell_type. \n Each cell type should contain at least N_celltype genes. \n Only provide the gene list. \n Do not provide list under input cell types. \n Ensure each gene is listed in a new line without any prefixes such as numbers.”

In this template, “Cell_type” and “N_celltype” can be replaced with the corresponding parameters.

The default GPT model used in tpGPT is gpt-4o. However, users can specify other models to generate the gene panel, such as gpt-4-turbo and gpt-3.5-turbo, within the tpGPT function.

#### Evaluation metrics

To evaluate the clustering performance, we calculated two metrics with known cell type labels.

The first metric is Jaccard index (JI). This metric is used to evaluate the similarity between the ground truth set and the set obtained by clustering. It is defined as the size of the intersection divided by the size of the union of the sample sets. Let TP, FN, and FP denote True Positives, False Negatives and False

Positives.The Jaccard Index is defined as follow 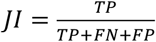.

The second metric is Adjusted Rand Index (ARI). Unlike the Jaccard Index, the ARI adjusts for the chance grouping of elements, providing a more rigorous assessment of clustering. It considers all pairs of samples and counts pairs that are assigned in the same or different clusters in the predicted and true clusters. The ARI is the corrected-for-chance version of the Rand index. Let *U* = {*u*_1_,…, *u*_*r*_} and V = {*v*_1_,…, *v*_*s*_} be two partitions on dataset X. And ARI is defined as,

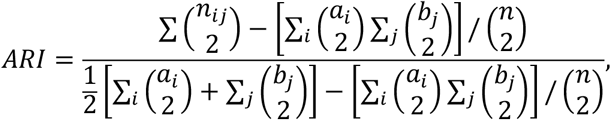

where 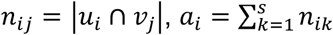 and, 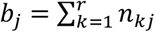.

Data processing and Seurat object: To proceed the downstream analysis such as UMAP clustering, we transfer the original scRNA-data and imputed data into a Seurat object with meta information ^19^. The UMI count data is normalized with Normalization function in Seurat package.

UMAP visualization with Seurat: To visualize the high-dimensional scRNA-seq data post-imputation, we utilized the Uniform Manifold Approximation and Projection (UMAP) technique implemented in the Seurat package. UMAP is a non-linear dimensionality reduction method that helps to uncover the intrinsic structure and relationships within complex datasets. Each dataset is first scaled to normalize the gene expression measurements, ensuring that variable genes are appropriately weighted. The scaled data is then subjected to principal component analysis (PCA) to reduce the dimensions of the dataset to the principal components that capture the most variation. In our study, we use the first 50 PCAs for analysis.

Cell type annotation in the normalized HNSCC data: Cell types within the normalized HNSCC data were predicted using the SingleR algorithm, a methodology that leverages reference data to assign cell type labels to individual cells in the experimental dataset. We search with the default references used in singleR package with “Human” species. The results of the cell type annotation are presented with the reference of BLUEPRINT which contains the most cell types in the normalized HNSCC dataset. We also utilized the AUCell package to verify cell types within this dataset. AUCell calculates the Area Under the Curve (AUC) to assess the activity of predefined gene sets within individual cells. We constructed marker gene sets for each cell type by selecting the top five highly expressed genes characteristic of the corresponding cell type. These marker gene sets were then used to build gene rankings for each cell based on their expression profiles. By calculating the AUC scores for these gene sets, we were able to determine the relative activity of the marker genes in each cell. Cells exhibiting high AUC scores for a particular gene set were identified as belonging to the corresponding cell type.

Cell type annotation in the HNSCC count data: we employed Azimuth cell type annotation application built with Seurat package (https://satijalab.org/azimuth/). The publicly available peripheral blood mononuclear cell (PBMC) reference data were selected and incorporated as the reference data in “RunAzimuth” function. The choice of the PBMC dataset was guided by its alignment with the expected cell types inherent in the HNSCC count data.

The annotation accuracy is calculated using the F1 score which is defined as: 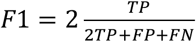.

TP, FN, and FP denote True Positives, False Negatives, and False Positives, respectively, as defined in the calculation of the Jaccard Index.

#### Performance benchmarking against existing methods

We benchmark the performance of SmartImpute with other imputation frameworks as follows:

SAVER (v1.1.3): Single-cell analysis via expression recovery (SAVER) is an expression recovery method using a Bayesian Framework. SAVER assumes a Poisson-Gamma mixture model to estimate the posterior distribution of the gene expression levels. SAVER employs a LASSO penalty which has the similar functions to the penalization we used in the loss function *L*_2_. We used the SAVER R package with default parameters, as detailed in the SAVER documentation (https://github.com/mohuangx/SAVER).

DCA: The Deep Count Autoencoder (DCA) framework uses a zero inflated negative binomial noise model to impute the scRNA-seq data. By incorporating the ZNIB distribution model with an autoencoder framework, DCA can capture the sparsity and overdispersion of the count data. We followed the default setting with learning rate 0.001, batch size 32 and a latent representation space of 32 dimensions. All other hidden layers are 64 dimensions. (https://github.com/theislab/dca).

DeepImpute: DeepImpute employs a data driven targeted imputation approach with deep neural network. The first step is using the variance over mean ration with default value 0.5 to select the target genes.

DeepImpute splits the genes into random subsets with each containing S genes. We follow the default setting of S=512, aligning with the number of target genes used in SmartImpute. We set the learning rate to 0.0001 and a batch size 64 as suggested in default settings. DeepImpute contains a dropout layer and we set the dropout rate as 20% as suggested by the author. (https://github.com/lanagarmire/DeepImpute).

The computation time for each method was evaluated using a subset of the normalized HNSCC data on a 3.3 GHz 12-Core Xeon Processor machine. We calculated the average running time five scenarios involving different sizes of gene subsets. In each subset, we included the targeted panel genes of SmartImpute (size = 580) and randomly added additional genes, resulting in total gene numbers for each scenario of 1,000, 5,000, 10,000, 15,000, and 18,241. All cells (total = 5,357) were used in each scenario. Each method was run five times to obtain the average computation time. All imputation results were converted into Seurat objects for subsequent UMAP clustering.

## Acknowledgements

This work has been supported in part by a National Institutes of Health grant [R01DE030493 to X.W]; and Biostatistics and Bioinformatics Shared Resources at the H. Lee Moffitt Cancer Center & Research Institute, an NCI-designed Comprehensive Cancer Center (P30-CA076292).

