## Supplementary Figures for "SmartImpute: A Targeted Imputation Framework for Single-cell Transcriptome Data"

Supplementary Figure 1

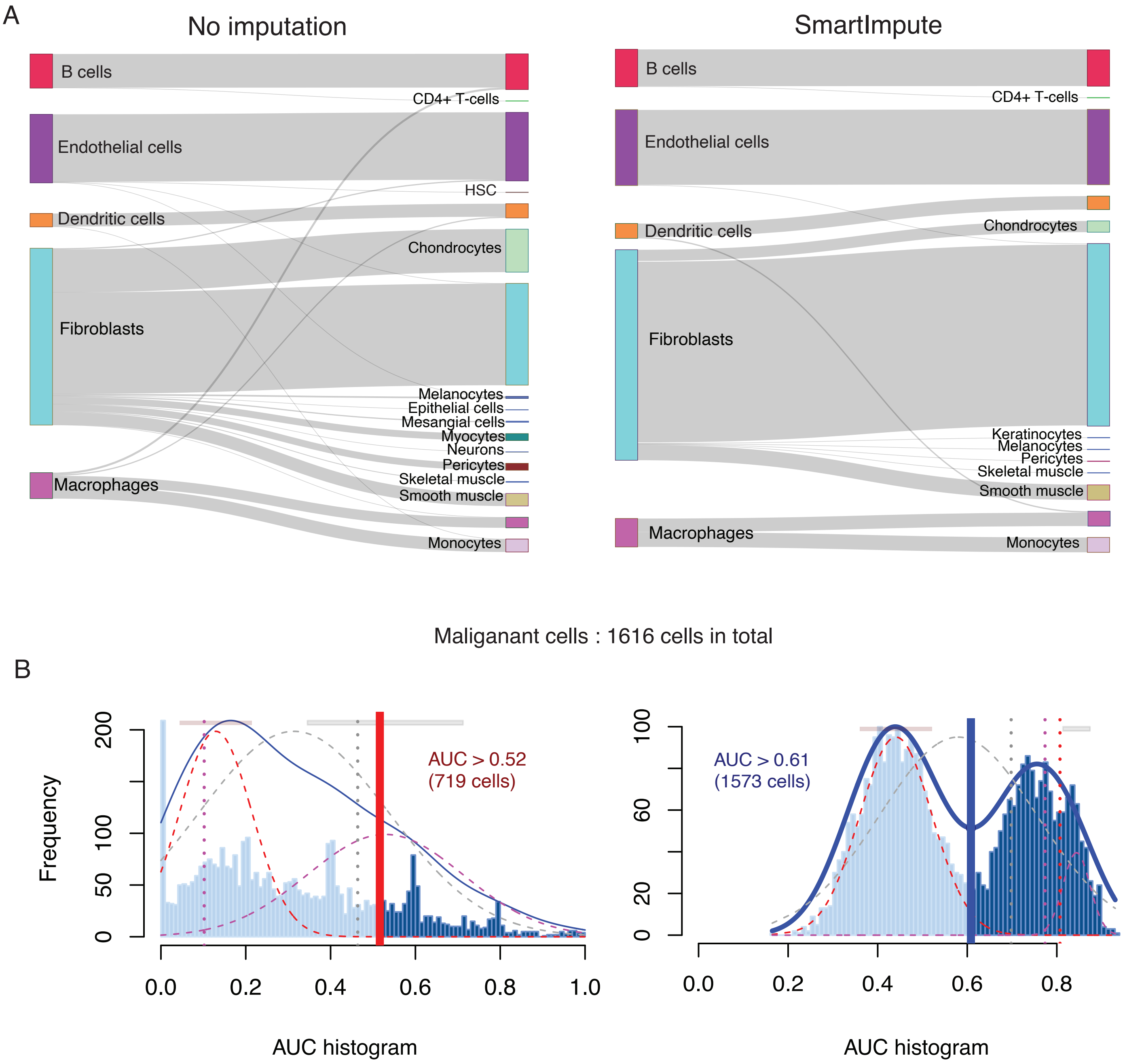


**Supplementary Figure 1.** SmartImpute performance in HNSCC normalized data. (A) Cell type annotation results with singleR: no imputation (left) and SmartImpute (right). (B) AUCell annotation results in malignant cells: no imputation (left) and SmartImpute (right)

Supplementary Figure 2

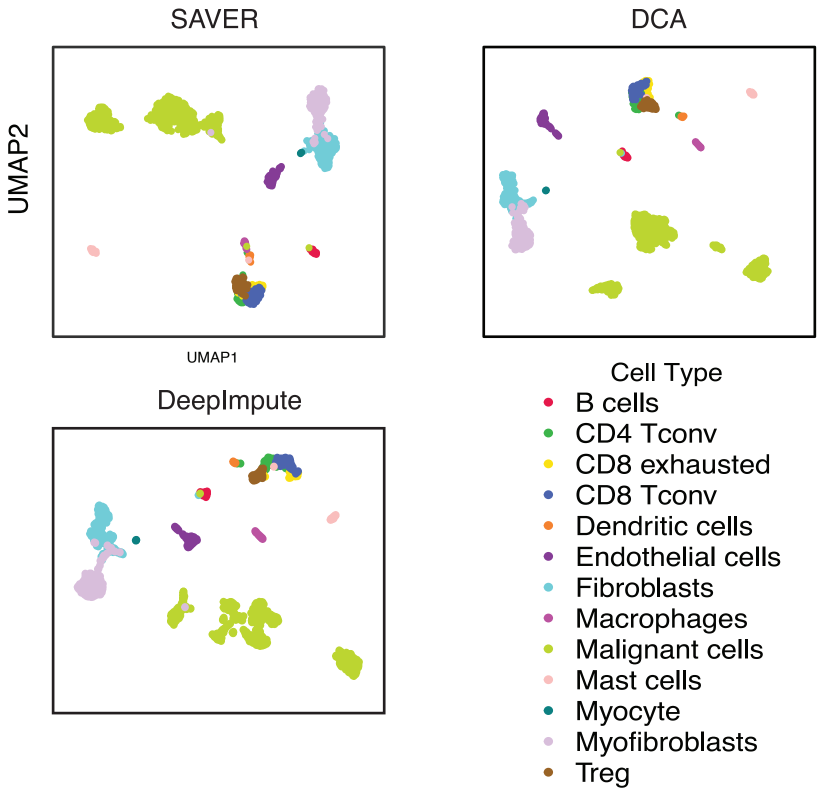


**Supplementary Figure 2.** UMAP clustering results for state-of-the-art imputation methods in HNSCC normalized data: SAVER, DCA and DeepImpute

Supplementary Figure 3

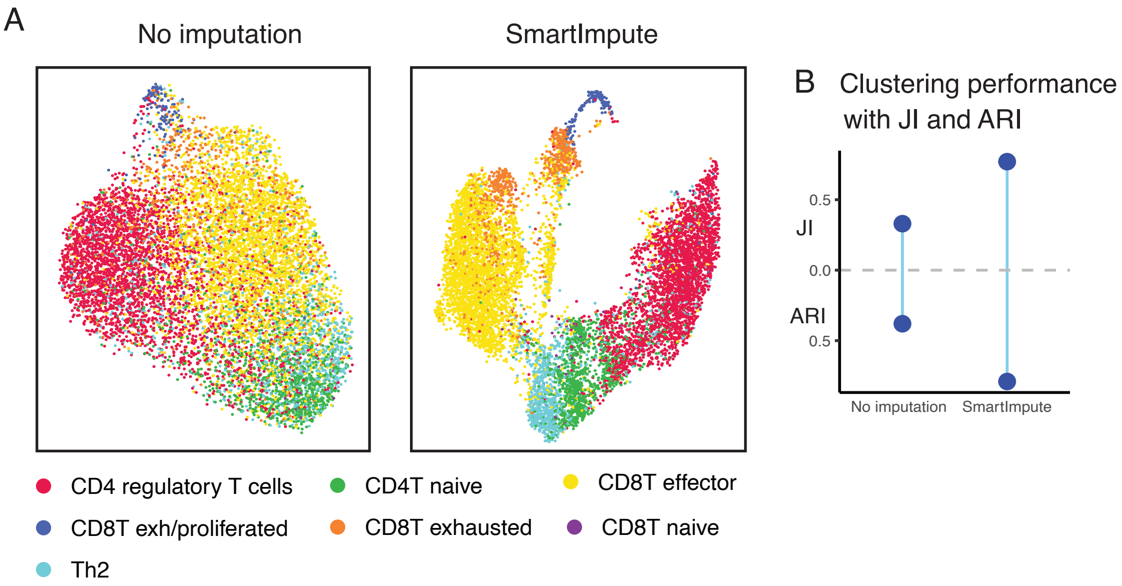


**Supplementary Figure 3.** Performance of SmartImpute in T cell subtypes. (A) UMAP clustering: no imputation (left) and SmartImpute (right). (B) Clustering measurement with Jaccard Index (JI) and Adjusted Rand Index (ARI)

Supplementary Figure 4

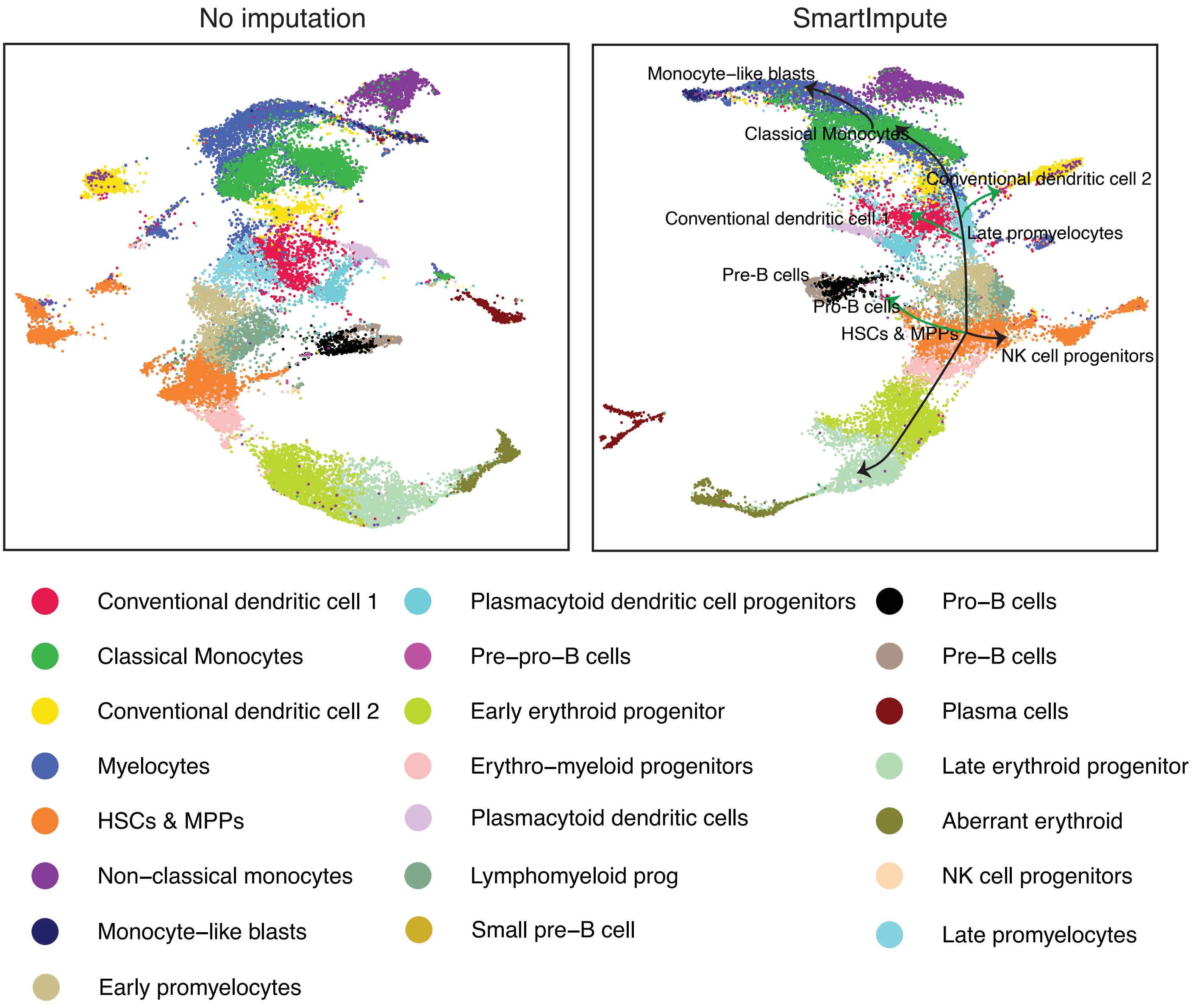


**Supplementary Figure 4.** Human bone marrow celluar trajectories with SmartImpute
